# Natural variation in male frequency fails to predict inbreeding responses in *Caenorhabditis elegans*

**DOI:** 10.64898/2026.05.07.723510

**Authors:** Jasbelle Sosa, Steve Abraham, Gabriel A. Blanco, Jessica Olivera, Isabella Alonso, Janna L Fierst, Rohit Kapila

## Abstract

In androdioecious species like *Caenorhabditis elegans*, where the primary mode of reproduction is self-fertilization, the evolutionary role of males has long puzzled biologists. One proposed benefit of males is the potential to escape inbreeding depression. We tested this by enforcing seven generations of inbreeding across nine *C. elegans* strains differing in baseline male frequency and measuring competitive relative fitness before and after inbreeding. We then relaxed inbreeding for four generations to assess recovery. We predicted that strains with higher male frequency, and greater opportunity for outcrossing, would exhibit faster recovery once inbreeding was relaxed. Strains varied substantially in their responses with most showing significant fitness declines and partial recovery but neither the magnitude of inbreeding depression nor the extent of recovery correlated with male frequency. These results show that male frequency is a poor predictor of inbreeding responses and does not reliably reflect realized outcrossing or its fitness consequences.

## Introduction

Despite its inherent disadvantages, such as lower reproductive output, lower population expansion rates, and associated costs like predation risk, sexual reproduction remains the dominant mode in most species (1). One of the primary arguments favouring sexual reproduction is its potential to enhance a species’ long-term evolutionary adaptability by introducing genetic variation (2). Androdioecious species like *C. elegans*, where self-fertilizing hermaphrodites coexist with rare males, pose a unique puzzle to this paradigm. Despite retaining the machinery for sexual reproduction, such populations rely predominantly on selfing which should erode the genetic benefits of sex while retaining demographic costs. Understanding why males persist at low frequencies in these systems offers key insights into the evolutionary maintenance of sex.

One proposed explanation for the persistence of males in androdioecious species is that facultative outcrossing may help populations escape inbreeding depression by restoring heterozygosity and purging deleterious mutations (3–5). A common assumption is that higher male frequency increases opportunities for outcrossing and thus may mitigate inbreeding depression. Across studies, reported male frequencies in *C. elegans* vary from near-zero spontaneous male production in many wild strains (∼0.05–0.12%) (6) to much higher levels in some natural isolates or experimentally maintained populations, reaching ∼35–40% under laboratory conditions (7). Males in these populations may confer increased resilience to inbreeding depression through increased heterozygosity, exposing deleterious variation and allowing it to be purged from populations. The maintenance of segregating variation in these populations may also allow them to recover more quickly from inbreeding.

Alternately, outcrossing can disrupt coadapted gene complexes and reduce fitness (8–11). Previous studies with the N2 lab adapted strain of *C. elegans* found that populations suffer from outbreeding depression (9). Highly heterozygous outcrossing *Caenorhabditis* species exhibit high levels of inbreeding depression (9) and struggle to recover lost fitness or purge genetic load (12). These factors raise a key question: can variation in male frequency predict how populations respond to inbreeding and recover from it?

To address this, we used nine genetically distinct strains of self-fertilising *Caenorhabditis* elegans with natural variation for male frequency. Each strain was subjected to seven generations of enforced inbreeding, followed by four generations of recovery under relaxed inbreeding. Previous work in *C. elegans* has shown that male frequency varies across strains and environments (13–15), making it a powerful system to test these ideas. However, it remains unclear whether natural variation in male frequency is a reliable predictor of inbreeding depression and recovery. We hypothesized that strains with naturally higher male frequencies would exhibit: 1) Reduced fitness loss under enforced inbreeding; and 2) Improved recovery after inbreeding. Under these scenarios, historical opportunities for outcrossing have both maintained higher heterozygosity and reduced the accumulation or expression of recessive deleterious alleles. Alternatively, male frequency may itself correlate with other underlying genetic properties affecting resilience to inbreeding depression.

## Materials and methods

### Strains and maintenance

We worked with nine focal strains of *C. elegans*: JU345, MY16, N2, MDX44, JU311, PX176, ECA36, AB1, and CB4856, and one competitor ST2. ST2 carries a GFP marker and each focal strain was measured against the same ST2 background to provide a standardized competitive reference. Thus, any fitness effect associated with the ST2 genetic background or GFP marker is expected to be constant across assays, and observed differences in relative fitness primarily reflect intrinsic variation among focal strains and experimental treatments.

Each focal strain was maintained as three replicate subpopulations throughout the experiment. All strains were obtained from the Caenorhabditis Genetics Center (CGC) and stored at −80 °C. Frozen strains were thawed onto 75 mm × 13 mm Nematode Growth Medium (NGM) plates seeded with E. coli OP50. After one generation of recovery (∼3.5 days at 20 °C), each focal strain population was divided into three replicate subpopulations, which were maintained separately as experimental blocks throughout the experiment. The common competitor strain ST2 was maintained as a single laboratory stock without replication. Prior to each assay, worms from each separately maintained block were synchronized by bleaching to obtain age-matched L1 larvae. Briefly, gravid adults from each block were lysed in hypochlorite solution, and the recovered embryos were allowed to hatch on 75 mm × 13 mm NGM plates seeded with E. coli OP50 to ensure developmental synchronization (16).

### Enforced inbreeding protocol

To enforce inbreeding, a single hermaphrodite carrying visible eggs was randomly selected every seven days and transferred to a fresh NGM plate. This single-worm transfer design reduces effective population size and is expected to increase homozygosity through repeated selfing. The transfer was repeated for seven consecutive weeks for each focal strain, effectively propagating each line through repeated self-fertilization. All replicate blocks were maintained independently throughout the inbreeding process. This approach is widely used to generate inbred lines in *C. elegans* (17). After 7 weeks of enforced inbreeding all the worms were stored at −80 °C until further use. Because enforced inbreeding prevented outcrossing, this phase tests whether baseline male frequency predicts resilience to inbreeding rather than realized outcrossing.

### Recovery from inbreeding protocol

Frozen strains were thawed onto 75 mm × 13 mm NGM plates seeded with *E. coli* OP50. For the next four weeks, large chunks of agar containing worms were transferred weekly to fresh NGM plates seeded with E. coli OP50. Care was taken to ensure that each transfer included >1000 mixed-stage worms to minimize sampling effects and demographic bottlenecks. This expansion phase allows population growth and selection from residual genetic variation and outcrossing if males are present. Because male frequency was not quantified during recovery, the recovery metric reflects overall fitness rebound and may include the composite effects of selection, recombination and outcrossing.

### Experimental assays

All the assays were conducted across three independent blocks, with a target of 10–15 individuals per strain per block. Prior to each assay, populations were bleach-synchronized as described above to obtain age-matched cohorts and minimize variation due to developmental stage. After excluding individuals that died or yielded incomplete data, final sample sizes ranged from approximately 7–14 individuals per block per strain.

### Male frequency assay protocol

Total progeny produced by each worm over four days were sexed to estimate male frequency. For each strain and replicate, 10–11 hermaphrodites were individually assayed. A single late-L4 hermaphrodite was placed on an NGM plate seeded with *E. coli* OP50 and transferred to a fresh plate every 24 hours for four consecutive days. Progeny from each plate were allowed to mature to adulthood and were then sexed and counted. If a worm died during the transfer series, its data were excluded from analysis. Male frequency was calculated as the ratio of the total number of males produced across four days to the total number of adult progeny scored.

### Competitive relative fitness assay

Five late-L4 hermaphrodites from the focal strain and two late-L4 hermaphrodites from a common GFP-marked competitor strain (ST2) were placed together on 75 mm × 13 mm NGM plates seeded with E. coli OP50. Mixed populations were allowed to grow and reproduce on the same plate for eight days (∼two generations).

On day 8, worms were washed off the plates using M9 buffer and suspended in M9 containing 5 mM levamisole to immobilize them. A 5 µL drop of the suspension was placed on a glass slide and covered with a coverslip. Each sample was imaged twice using a fluorescence microscope: once in bright-field mode to capture all worms, and once under GFP excitation to visualize only the GFP-tagged ST2 competitor worms. The number of GFP-positive competitor worms (G) and total worms (NG) was counted from these images. The number of focal worms was calculated as (NG − G). Relative fitness (RF) of the focal strain was estimated from the change in focal-to-competitor ratio during the competition assay as:

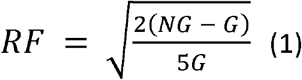

where 5/2 represents the initial focal-to-competitor ratio and the square root accounts for approximately two generations of competition. Values greater than 1 indicate that the focal strain increased in frequency relative to the competitor, whereas values less than 1 indicate a competitive disadvantage. Image acquisition and manual counting of GFP-marked and non-fluorescent worms followed established image-based competitive fitness procedures (14,17,18), adapted here for plate-based assays. This metric reflects relative competitive reproductive output and is monotonic with standard measures of relative fitness.

### Developmental time

To assess whether strain differences in relative fitness could be confounded by variation in life-history timing, we measured developmental time (L1 to first reproduction) under common-garden conditions. Because experimental populations were maintained on plates for fixed durations (∼8 days), substantial differences in developmental rate could lead to unequal numbers of reproductive cycles across strains. All the replicates of all the strains were bleached for age synchronisation, eggs were left suspended overnight in M9 solution for further age synchronisation. One L1 was placed on a plate and was observed every hour starting 45 hours post bleaching until the first egg was observed on the same plate. 11 L1s per replicate per strain replicate were observed.

## Statistical analysis

To test how inbreeding and recovery affected relative fitness, we fitted a linear mixed-effects model using the *glmm*TMB package in R (19), with Status (Before inbreeding, After inbreeding, After recovery), Strain, and their interaction as fixed effects, and Block as a random intercept to account for experimental variation:

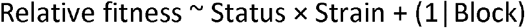

Model significance was evaluated using Type III Wald chi-square tests using the car::Anova() function (20). Post-hoc pairwise contrasts were performed using the *emmeans* package (21) with Tukey adjustment to control for multiple comparisons among the three contrasts per strain.

Model diagnostics were assessed using *DHARMa(*22). While the initial model showed deviations from residual uniformity (Kolmogorov–Smirnov test, p < 0.001) and an elevated frequency of outliers (p < 0.001), log-transformation of relative fitness resolved these issues (KS test, p = 0.31; outlier test, p = 0.66). Importantly, the qualitative conclusions were unchanged under the alternative specification, with Type III Wald chi-square tests confirming significant effects of Status, Strain, and their interaction in the log-transformed model.

From the estimated marginal means, we derived strain-specific indices of inbreeding depression and recovery to quantify proportional changes in relative fitness across phases. Inbreeding depression (ID) was calculated as (Before inbreeding − After inbreeding) / Before inbreeding, representing the fractional decline in relative fitness due to inbreeding. Recovery (IR) was calculated as (After recovery − After inbreeding) / Before inbreeding, representing the proportional change in relative fitness following the relaxation of inbreeding relative to the inbred state. Positive IR values indicate improvement (fitness rebound), values near zero indicate no change, and negative values indicate that mean relative fitness after recovery was lower than during inbreeding. We then tested whether natural male frequency predicted either of these responses using Spearman’s rank correlations and simple linear regressions between male frequency and both ID and IR. To assess whether variation in developmental time influenced strain-level estimates of inbreeding depression and recovery, we fitted additional linear models including mean strain developmental time as a predictor, both alone and together with mean male frequency. Collinearity between predictors was assessed using Spearman’s rank correlation and variance inflation factors (VIF).

## Results

Male frequency varied from 0% to 7.68% across strains (mean ± SD = 1.40 ± 2.46%), with high among-strain variability (Table S1). The mixed-effects model revealed significant effects of Status (χ^2^ = 7.10, p = 0.0288) and Strain (χ^2^ = 53.38, p = 9.09 × 10^−9^), as well as a significant Status × Strain interaction (χ^2^ = 66.6, p = 3.82 × 10^−8^; Table 1). Relative fitness generally declined following inbreeding, but both the magnitude of decline and the extent of recovery varied substantially among strains. Recovery after four generations of relaxed inbreeding was incomplete and highly variable (Fig. 1; Tables 1; Table S2).

**Table 1.** Type III Wald χ^2^ tests from the mixed-effects model examining the effects of inbreeding status, strain, and their interaction on relative fitness.

| *Effect* | $\chi^2$ | *df* | *p value* |
| --- | --- | --- | --- |
| (Intercept) | 168.63 | 1 | $< 2.2 \times 10^{-16}$ |
| Status | 7.10 | 2 | 0.0288* |
| Strain | 53.38 | 8 | $9.09 \times 10^{-9***}$ |
| Status × Strain | 66.66 | 16 | $3.82 \times 10^{-8***}$ |

**Figure 1.**
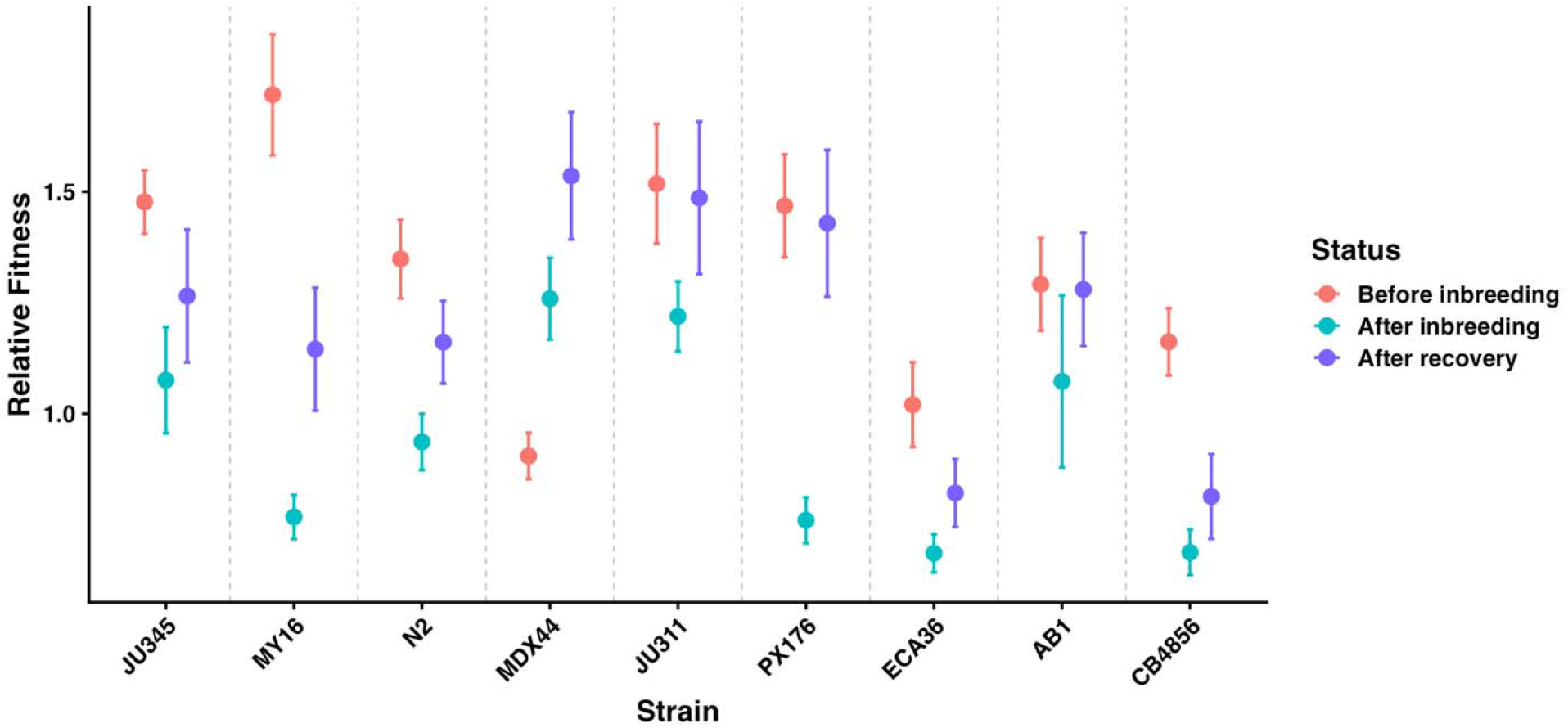
Mean ± SE of relative fitness across nine *C. elegans* strains before inbreeding, after seven generations of enforced inbreeding, and after four generations of recovery. Strains are ordered by increasing male frequency (left to right). Colours indicate experimental stage. Values of relative fitness greater than 1 indicate that the focal strain outcompeted the GFP-marked competitor, whereas values less than 1 indicate a competitive disadvantage.

### Strain-specific contrasts revealed distinct responses to inbreeding and recovery

While several strains exhibited significant declines in relative fitness following inbreeding, others showed little or no evidence of inbreeding depression, and one strain (MDX44) exhibited increased fitness under inbreeding. Recovery patterns were similarly variable. For example, MY16 and PX176 showed significant recovery following relaxation of inbreeding, although MY16 did not return to pre-inbreeding fitness levels, whereas CB4856 remained depressed after recovery. In contrast, MDX44 exhibited consistently high fitness across phases, exceeding its pre-inbreeding level after recovery. Together, these results indicate that both susceptibility to inbreeding depression and the capacity for recovery depend strongly on genetic background (Table S2; Fig. 1). Such variability is consistent with recent work in outcrossing *Caenorhabditis* showing that the magnitude of inbreeding depression can differ substantially across species and populations (23).

To test whether among-strain variation could be explained by differences in male frequency we examined the relationship between population male frequency and inbreeding response. Despite substantial variation in relative fitness across strains, male frequency did not predict either the magnitude of inbreeding depression or the extent of recovery (Fig. 2; Table S4). Neither response was significantly associated with male frequency in rank-based correlation analyses (Spearman: ID, ρ = −0.042, p = 0.915; IR, ρ = −0.176, p = 0.651). Linear models also showed weak, non-significant effects of male frequency on both inbreeding depression (slope = 1.70 ± 4.02, p = 0.685) and recovery (slope = −1.42 ± 1.52, p = 0.383), with 95% confidence intervals overlapping zero (Table S4).

**Figure 2.**
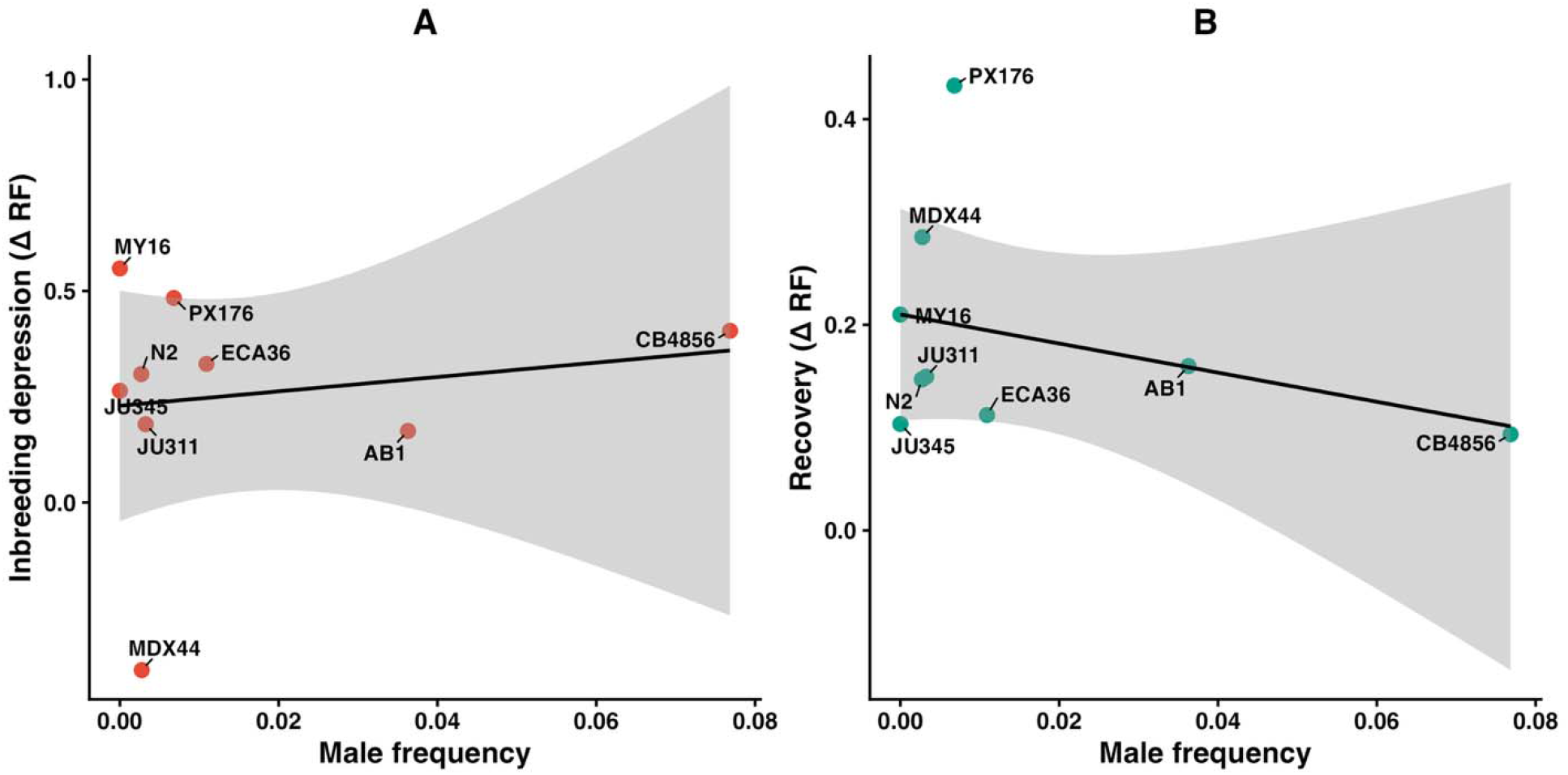
Relationship between natural male frequency and (A) inbreeding depression (ID) and (B) recovery (IR) across nine *C. elegans* strains. Inbreeding depression (ID) was calculated as the proportional decline in relative fitness from before inbreeding to after inbreeding, and recovery (IR) as the proportional change in relative fitness from after inbreeding to after recovery. Points represent strain-level estimates derived from model estimated marginal means. Solid lines indicate linear regression fits, with shaded areas representing 95% confidence intervals.

Further, to directly compare explanatory power, we quantified variance explained by genetic background and by male frequency. Genetic background explained substantially more variation in relative fitness responses (R^2^ = 0.216, adjusted R^2^ = 0.187) than male frequency explained in either inbreeding depression (R^2^ = 0.025, adjusted R^2^ = −0.114) or recovery (R^2^ = 0.110, adjusted R^2^ = −0.017), indicating that responses were strongly strain-dependent, whereas male frequency explained little variation (Fig. S1).

Developmental time varied from 63.0 to 70.2 hours across strains (mean ± SD = 66.6 ± 2.3 hours; Fig. S2; Table S5) and was not correlated with male frequency across strains (Spearman’s ρ = −0.10, p = 0.797; VIF = 1.00), indicating minimal collinearity between predictors. Developmental time showed a suggestive negative association with inbreeding depression when modeled alone (β = −0.078 ± 0.035, p = 0.061), with faster-developing strains tending to exhibit greater declines in relative fitness (Fig. S3). However, in a model including both male frequency and developmental time, male frequency remained non-significant (β = 1.52 ± 3.31, 95% CI: −6.57 to 9.61, p = 0.662), whereas developmental time showed only a suggestive negative trend (β = −0.077 ± 0.037, 95% CI: −0.168 to 0.013, p = 0.082). For recovery, neither male frequency (β = −1.43 ± 1.62, 95% CI: −5.39 to 2.52, p = 0.409) nor developmental time (β = −0.008 ± 0.018, 95% CI: −0.053 to 0.036, p = 0.665) were significantly different between populations (Table S6).

## Discussion

There is a long-standing debate about the evolutionary significance of males in *C. elegans*, a predominantly selfing species in which males are maintained at low frequencies (24,25). One proposed advantage is that males mitigate the fitness costs of inbreeding by enabling outcrossing, thereby restoring heterozygosity and facilitating recombination, as suggested for other androdioecious systems(9,26,27). We found no evidence that natural variation in male frequency predicts either the magnitude of inbreeding depression or the extent of recovery across *C. elegans* strains. Despite substantial among-strain variation in relative fitness responses, both inbreeding depression and recovery were more strongly associated with genetic background than male frequency. These findings suggest that male frequency is a poor proxy for the capacity of populations to mitigate inbreeding depression. Thus, neither of our initial predictions, that higher male frequency reduces inbreeding depression or accelerates recovery, were supported. More broadly, our results challenge the assumption that baseline levels of males reliably reflect realized outcrossing or its consequences for fitness recovery.

Across strains, the effects of inbreeding and recovery were highly variable, indicating that inbreeding resilience is strongly dependent on genetic background. While some strains (e.g., N2 and PX176) exhibited partial recovery following relaxed inbreeding, others such as JU345 and CB4856 remained depressed despite representing opposite extremes of male frequency. This mismatch between mating system variation and inbreeding response suggests that baseline male frequency may not reliably capture realized outcrossing or the potential for genetic rescue. Comparative work across *Caenorhabditis* further suggests that mating system may shape the distribution of genetic load, with obligate outcrossers often harboring greater concealed deleterious variation, whereas predominantly selfing species tend to exhibit reduced inbreeding depression through purging (23). Instead, variation in responses may reflect lineage-specific evolutionary histories, including differences in mutation load, past selection, recombination landscapes, or developmental traits. The exceptional robustness of MDX44, which maintained or even improved reproductive output under inbreeding, may indicate prior purging of deleterious alleles or other compensatory genetic mechanisms. An earlier study found a similar pattern of outbreeding depression in the lab-adapted N2 model strain of *C. elegans* (9). Prior work with C. remanei identified a complex relationship between inbreeding and fitness recovery in this highly diverse obligate outbreeding species (12). This and other work indicates the genetic architecture of segregating variation does not directly predict how a species will respond to either inbreeding or recovery.

Our inbreeding design involved small effective population sizes, under which genetic drift may have contributed to stochastic differences in the fixation or purging of deleterious alleles during inbreeding. Because enforced inbreeding prevented outcrossing during the inbreeding phase, our results do not directly test whether realized outcrossing reduces inbreeding depression. Instead, they test whether naturally varying male frequency predicts strain-level resilience to inbreeding, potentially through correlated genetic or evolutionary-history differences. However, similar designs are widely used in *C. elegans* studies of inbreeding and mutation accumulation, allowing direct comparison with previous work(17). In addition, our fitness assay measured competitive reproductive output over a fixed interval, capturing only one component of organismal performance. Differences in intrinsic life-history traits, such as developmental time and reproductive timing, may therefore influence apparent fitness responses in this assay. Although developmental time showed a weak negative association with inbreeding depression (Fig. S2, S3), this relationship was only suggestive, did not extend to recovery, and did not alter the conclusion that baseline male frequency was not predictive of either inbreeding depression or recovery. Together, these findings challenge the assumption that greater potential for outcrossing necessarily translates into increased resilience to inbreeding(28–30).

A likely explanation for this pattern is that male frequency does not necessarily reflect realized outcrossing rates or the effective recombination occurring within a population. In *C. elegans* a single male can mate with multiple hermaphrodites, potentially limiting the marginal benefit of additional males. Moreover, strain-specific differences in mating efficiency, sperm precedence, and hermaphrodite receptivity may decouple male abundance from its functional contribution to genetic mixing. Behavioural differences among strains may further influence encounter rates and mating success. *C. elegans* males are notoriously poor at mating, both finding hermaphrodites and accurately placing sperm(31–33). As inbreeding decreases population-fitness the mating competence of *C. elegans* males may be further eroding. Thus, males may be present but unable to successfully mate at a level necessary to rescue the population from inbreeding depression.

Across the broader *C. elegans* literature, our results fit a recurring pattern: male frequency and outcrossing potential are highly context- and background-dependent, rather than simple predictors of resilience. Under standard laboratory conditions, males are often selected against and tend to decline toward low frequencies (14), and even under elevated mutational pressure, transient increases in male frequency do not necessarily lead to the invasion or maintenance of obligate outcrossing(34). Similarly, the expression of partially recessive deleterious alleles can reduce the cost of males and maintain them at higher frequencies, but these effects vary among genetic backgrounds (35). Comparative and experimental evolution studies likewise suggest that mating-system dynamics, mutation responses, and the maintenance of genetic load depend strongly on strain-specific history and the balance between selfing and outcrossing (7,14).

Classical theory predicts that recombination and male function should facilitate the purging of deleterious mutations and accelerate recovery from inbreeding(30). Our results indicate that this advantage does not scale with standing male frequency under benign laboratory conditions. Because male frequency may not reflect realized outcrossing, particularly under environmental stress, where mating behaviour and sex ratios can shift, standing male frequency alone may be a poor proxy for mutational robustness. Our results show that standing male frequency alone does not predict inbreeding responses, highlighting the need to measure realized outcrossing directly when testing the evolutionary role of sex. More broadly, similar dynamics may occur in other androdioecious systems, where male abundance may not reliably predict realized outcrossing or resilience to inbreeding.

## Acknowledgments

JLF, RK, JS and GAB were supported by NIGMS award R35147245. RK was also supported by a Postdoctoral Fellowship from the Alliance of Hispanic Serving Research Universities.

