## Supplementary data for "Natural variation in male frequency fails to predict inbreeding responses in *Caenorhabditis elegans*"

Table S1. Strain-level mean male frequency estimates. Values represent the proportion of male progeny produced per strain, calculated across individual hermaphrodite assays. Standard deviation (SD) and standard error (SE) are shown to indicate variability among replicates. The competitor strain ST2 is included for completeness but was excluded from statistical analyses in the main text.

| **Strain** | **Mean frequency** | **SD** | **SE** |
| --- | --- | --- | --- |
| CB4856 | 0.0768 | 0.1200 | 0.0203 |
| AB1 | 0.0363 | 0.0653 | 0.0112 |
| ECA36 | 0.0109 | 0.0450 | 0.00761 |
| PX176 | 0.00680 | 0.0292 | 0.00474 |
| JU311 | 0.00325 | 0.0142 | 0.00248 |
| MDX44 | 0.00276 | 0.0143 | 0.00224 |
| N2 | 0.00270 | 0.0106 | 0.00170 |
| ST2 | 0.000653 | 0.00296 | 0.000479 |
| JU345 | 0.0000 | 0.0000 | 0.0000 |
| MY16 | 0.0000 | 0.0000 | 0.0000 |

Table S2. Within-strain pairwise contrasts from estimated marginal means testing the effects of inbreeding and recovery on  relative fitness in nine *C. elegans* strains. Negative estimates indicate lower relative fitness compared to the reference phase, whereas positive estimates indicate higher relative fitness. P-values were Tukey-adjusted for multiple comparisons.

| **Strain** | **Contrast** | **Estimate** | **SE** | **df** | **t.ratio** | **p.value** |
| --- | --- | --- | --- | --- | --- | --- |
| JU345 | After inbreeding - Before inbreeding | -0.3901 | 0.1489 | 703 | -2.621 | 0.0243 |
| JU345 | After recovery - Before inbreeding | -0.2372 | 0.1490 | 703 | -1.592 | 0.2498 |
| JU345 | After recovery - After inbreeding | 0.1529 | 0.1571 | 703 | 0.973 | 0.5940 |
| MY16 | After inbreeding - Before inbreeding | -0.9511 | 0.1403 | 703 | -6.779 | <0.0001 |
| MY16 | After recovery - Before inbreeding | -0.5903 | 0.1527 | 703 | -3.866 | 0.0004 |
| MY16 | After recovery - After inbreeding | 0.3608 | 0.1527 | 703 | 2.363 | 0.0482 |
| N2 | After inbreeding - Before inbreeding | -0.4084 | 0.1415 | 703 | -2.886 | 0.0112 |
| N2 | After recovery - Before inbreeding | -0.2109 | 0.1519 | 703 | -1.389 | 0.3473 |
| N2 | After recovery - After inbreeding | 0.1975 | 0.1508 | 703 | 1.310 | 0.3900 |
| MDX44 | After inbreeding - Before inbreeding | 0.3571 | 0.1427 | 703 | 2.503 | 0.0336 |
| MDX44 | After recovery - Before inbreeding | 0.6142 | 0.1484 | 703 | 4.140 | 0.0001 |
| MDX44 | After recovery - After inbreeding | 0.2571 | 0.1484 | 703 | 1.732 | 0.1940 |
| JU311 | After inbreeding - Before inbreeding | -0.2808 | 0.1457 | 703 | -1.928 | 0.1316 |
| JU311 | After recovery - Before inbreeding | -0.0538 | 0.1489 | 703 | -0.361 | 0.9307 |
| JU311 | After recovery - After inbreeding | 0.2271 | 0.1541 | 703 | 1.473 | 0.3044 |
| PX176 | After inbreeding - Before inbreeding | -0.7082 | 0.1427 | 703 | -4.963 | <0.0001 |
| PX176 | After recovery - Before inbreeding | -0.0746 | 0.1666 | 703 | -0.448 | 0.8955 |
| PX176 | After recovery - After inbreeding | 0.6337 | 0.1666 | 703 | 3.804 | 0.0005 |
| ECA36 | After inbreeding - Before inbreeding | -0.3343 | 0.1415 | 703 | -2.362 | 0.0483 |
| ECA36 | After recovery - Before inbreeding | -0.2198 | 0.1519 | 703 | -1.447 | 0.3175 |
| ECA36 | After recovery - After inbreeding | 0.1145 | 0.1508 | 703 | 0.759 | 0.7281 |
| AB1 | After inbreeding - Before inbreeding | -0.2185 | 0.1403 | 703 | -1.558 | 0.2649 |
| AB1 | After recovery - Before inbreeding | -0.0121 | 0.1415 | 703 | -0.085 | 0.9960 |
| AB1 | After recovery - After inbreeding | 0.2065 | 0.1415 | 703 | 1.459 | 0.3114 |
| CB4856 | After inbreeding - Before inbreeding | -0.4700 | 0.1428 | 703 | -3.292 | 0.0030 |
| CB4856 | After recovery - Before inbreeding | -0.3617 | 0.1530 | 703 | -2.364 | 0.0481 |
| CB4856 | After recovery - After inbreeding | 0.1083 | 0.1507 | 703 | 0.719 | 0.7525 |

Table S3. Estimated marginal means (EMMs) of relative fitness for each *C. elegans* strain across experimental phases (Before inbreeding, After inbreeding, After recovery), obtained from the mixed-effects model. Values are shown as mean ± SE with 95% confidence intervals**.**

| **Strain** | **Status** | **Relative fitness (emmean ± SE)** | **95% CI (lower–upper)** |
| --- | --- | --- | --- |
| JU345 | Before inbreeding | 1.477 ± 0.114 | 1.254–1.700 |
| JU345 | After inbreeding | 1.087 ± 0.124 | 0.843–1.331 |
| JU345 | After recovery | 1.240 ± 0.124 | 0.996–1.484 |
| MY16 | Before inbreeding | 1.719 ± 0.114 | 1.495–1.942 |
| MY16 | After inbreeding | 0.768 ± 0.114 | 0.544–0.991 |
| MY16 | After recovery | 1.128 ± 0.129 | 0.876–1.381 |
| N2 | Before inbreeding | 1.345 ± 0.115 | 1.119–1.571 |
| N2 | After inbreeding | 0.937 ± 0.114 | 0.713–1.160 |
| N2 | After recovery | 1.134 ± 0.126 | 0.886–1.382 |
| MDX44 | Before inbreeding | 0.901 ± 0.115 | 0.675–1.128 |
| MDX44 | After inbreeding | 1.259 ± 0.115 | 1.032–1.485 |
| MDX44 | After recovery | 1.516 ± 0.122 | 1.276–1.756 |
| JU311 | Before inbreeding | 1.518 ± 0.114 | 1.295–1.742 |
| JU311 | After inbreeding | 1.237 ± 0.120 | 1.001–1.474 |
| JU311 | After recovery | 1.464 ± 0.124 | 1.221–1.708 |
| PX176 | Before inbreeding | 1.465 ± 0.115 | 1.239–1.691 |
| PX176 | After inbreeding | 0.757 ± 0.115 | 0.530–0.983 |
| PX176 | After recovery | 1.390 ± 0.144 | 1.108–1.673 |
| ECA36 | Before inbreeding | 1.020 ± 0.115 | 0.794–1.246 |
| ECA36 | After inbreeding | 0.686 ± 0.114 | 0.462–0.909 |
| ECA36 | After recovery | 0.800 ± 0.126 | 0.552–1.048 |
| AB1 | Before inbreeding | 1.291 ± 0.114 | 1.068–1.515 |
| AB1 | After inbreeding | 1.073 ± 0.114 | 0.850–1.296 |
| AB1 | After recovery | 1.279 ± 0.115 | 1.053–1.506 |
| CB4856 | Before inbreeding | 1.158 ± 0.117 | 0.928–1.387 |
| CB4856 | After inbreeding | 0.688 ± 0.114 | 0.464–0.911 |
| CB4856 | After recovery | 0.796 ± 0.126 | 0.548–1.044 |

Table S4. Linear model estimates for the effect of male frequency on strain-level indices of inbreeding depression (ID) and recovery (IR), calculated from estimated marginal means of relative fitness. Spearman rank correlations are reported in the main text.

| **Response** | **Predictor** | **Slope** | **SE** | **95% CI** | **p-value** |
| --- | --- | --- | --- | --- | --- |
| Inbreeding depression (ID) | Male frequency | 1.70 | 4.02 | −7.80 to 11.20 | 0.685 |
| Recovery (IR) | Male frequency | −1.42 | 1.52 | −5.01 to 2.18 | 0.383 |

Table S5. Strain-level developmental time estimates. Values represent mean developmental time (hours from late L4 to adulthood) for each strain, with standard deviation (SD) and standard error (SE) indicating variability among replicates. Minimum and maximum values show the observed range within each strain.

| **Strain** | **Mean (hrs)** | **SD (hrs)** | **SE (hrs)** | **Min (hrs)** | **Max (hrs)** |
| --- | --- | --- | --- | --- | --- |
| MY16 | 63.0 | 4.38 | 0.86 | 58.5 | 72.4 |
| PX176 | 64.5 | 5.19 | 1.11 | 58.3 | 74.3 |
| JU311 | 65.2 | 5.43 | 0.96 | 58.1 | 77.4 |
| ECA36 | 65.4 | 7.54 | 1.43 | 58.4 | 92.7 |
| CB4856 | 66.3 | 6.12 | 1.22 | 57.4 | 81.9 |
| AB1 | 66.4 | 5.92 | 1.06 | 58.7 | 81.9 |
| JU345 | 67.0 | 4.12 | 0.75 | 60.5 | 77.4 |
| ST2 | 69.0 | 5.16 | 0.99 | 62.3 | 87.2 |
| MDX44 | 69.4 | 8.06 | 1.34 | 60.2 | 92.4 |
| N2 | 70.2 | 4.39 | 0.94 | 62.3 | 77.4 |

Table S6. Effects of male frequency and developmental time on strain-level inbreeding depression (ID) and recovery (IR). Values are estimates from multiple linear models including both predictors.

| **Response** | **Predictor** | **Estimate** | **SE** | **95% CI** | **p-value** |
| --- | --- | --- | --- | --- | --- |
| ID | Male frequency | 1.52 | 3.31 | −6.57 to 9.61 | 0.662 |
| ID | Developmental time | −0.077 | 0.037 | −0.168 to 0.013 | 0.082 |
| IR | Male frequency | −1.43 | 1.62 | −5.39 to 2.52 | 0.409 |
| IR | Developmental time | −0.008 | 0.018 | −0.053 to 0.036 | 0.665 |

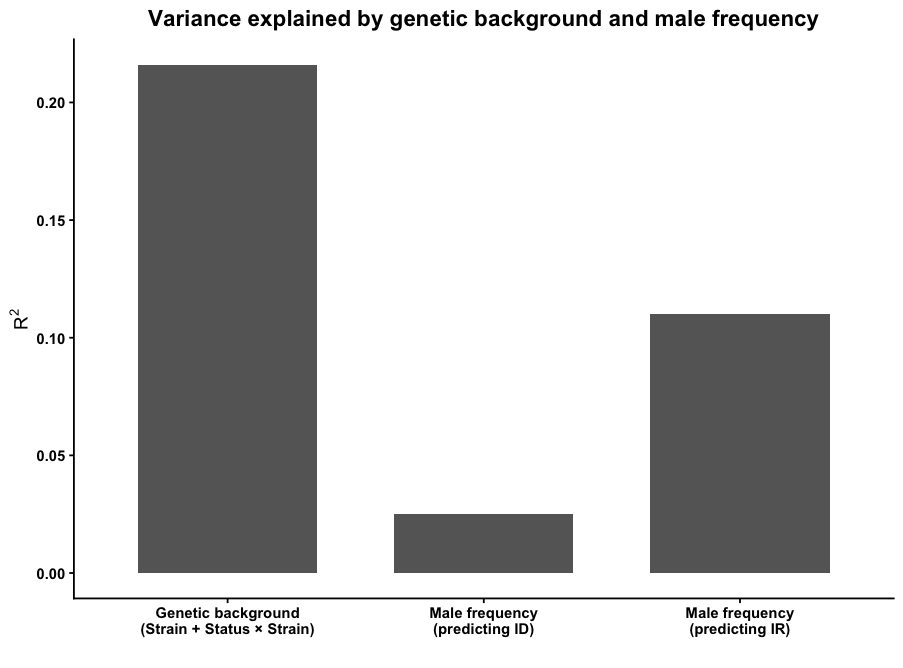

Figure 1: Genetic background explains substantially more variation in inbreeding responses than male frequency, which shows weak explanatory power for both inbreeding depression (ID) and recovery (IR).

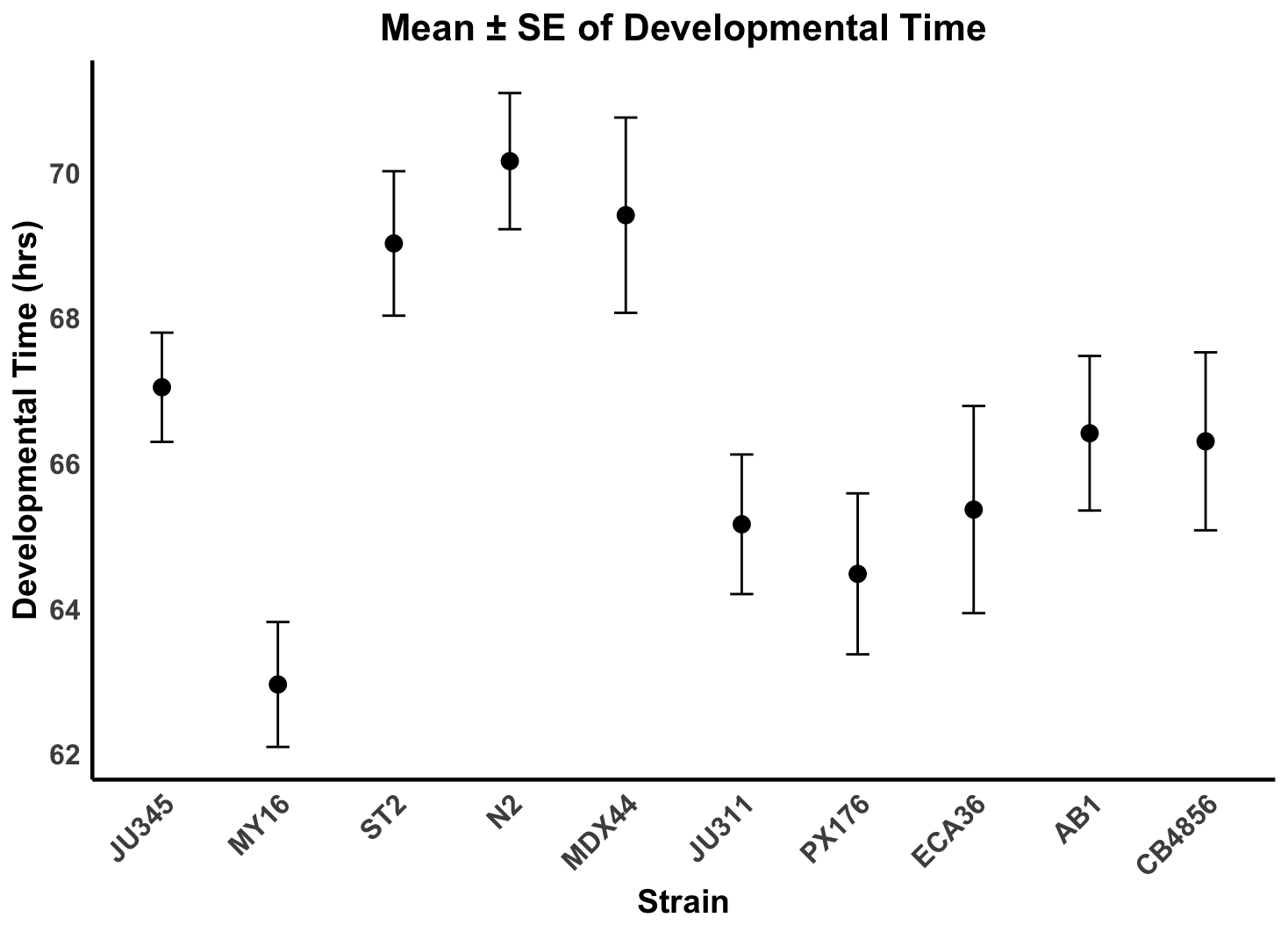

Figure S2: Mean ± SE of developmental time (hours from L1 to first reproduction) across nine C. elegans strains under common-garden conditions. Each point represents the strain mean based on replicate individuals. Developmental time differed significantly among strains, indicating inherent life-history variation beyond differences in male frequency.

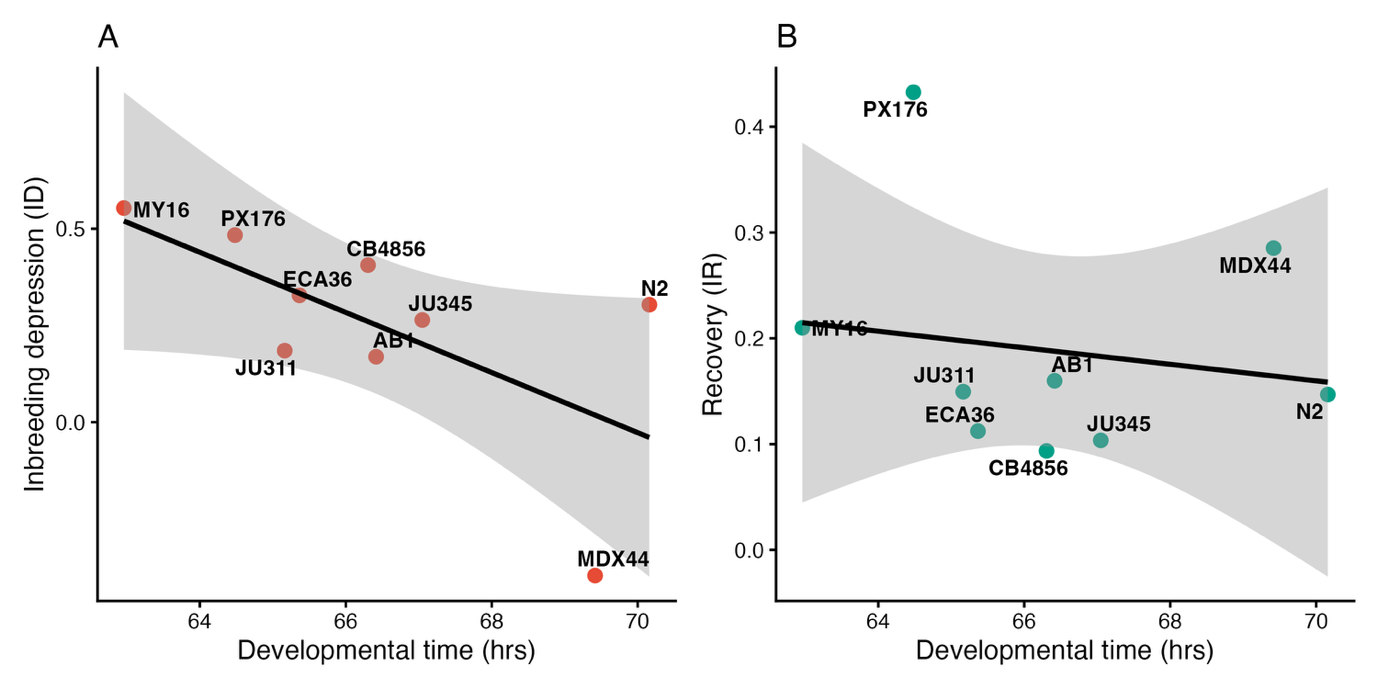

**Figure S3. Developmental time does not explain variation in inbreeding response.**

(A) Relationship between strain mean developmental time (L1 to first reproduction) and inbreeding depression (Δ relative fitness). (B) Relationship between developmental time and recovery following inbreeding. Points represent strain means; lines show linear model fits with 95% confidence intervals. Although a negative trend was observed between developmental time and inbreeding depression, this relationship was not statistically significant, and no association was detected for recovery. These results indicate that variation in relative fitness responses is unlikely to be explained by differences in developmental timing.
